# Vasopressin-Dependent β-Catenin Phosphorylation at Ser552 and Branching Structure of Mouse Collecting Duct System

**DOI:** 10.1101/2025.03.13.643123

**Authors:** Shuo-Ming Ou, Hiroaki Kikuchi, Euijung Park, Chin-Rang Yang, Viswanathan Raghuram, Shaza Khan, Adrian Rafael Murillo-de-Ozores, Lihe Chen, Chung-Lin Chou, Mark A. Knepper

## Abstract

**Background:** Phosphoproteomics studies in both cultured and native collecting duct (CD) cells showed that vasopressin strongly increases protein kinase A (PKA)-dependent phosphorylation of β-catenin at Ser552. Relatively little is known about the role of Ser552 phosphorylation.

**Methods:** To address the role of β-catenin Ser552 phosphorylation in the mature renal CD, we have inserted a Ser552Ala mutation in mice using CRISPR-Cas9.

**Results:** The mutation did not affect the renal abundance of the vasopressin-regulated water channel aquaporin-2 or urinary osmolality. However, the structure of the CD system was altered. Specifically, the cortical branching ratio (the number of nephrons that merge to form one cortical CD) was reduced from 6.18 ± 0.66 in control mice to 3.33 ± 0.82 in Ser552Ala mice. This was associated with a greater number of cortical and medullary CDs with smaller average diameter. The total number of nephrons (glomerular counts) was not different between wild-type and Ser552Ala mice (both ~13,500 per kidney). RNA-seq in microdissected cortical CDs of the mice revealed a highly significant enrichment of genes involved in regulation of mitosis and the cell cycle, along with decreases in mRNAs coding for two cyclin-dependent kinase inhibitor proteins, *Cdkn1b and Cdkn1c*. At the same time, there were no changes in abundances of major transporter mRNAs, indicative of sustained CD differentiation. A subset of cortical CD cells showed an increase in DNA content, consistent with G2/M cell-cycle arrest.

**Conclusions:** The observed structural changes in the collecting duct system of adult mice point to a role of vasopressin-mediated post-translational modification of β-catenin at Ser552 in collecting duct development, presumably PKA-mediated Ser552 phosphorylation. We speculate that vasopressin may act to slow or halt branching morphogenesis perinatally and may affect the collecting duct elongation process that normally produces the unbranched region of the CD system in the cortex and outer medulla.

**Key points:**

- Vasopressin regulates collecting duct (CD) transport by triggering phosphorylation of multiple proteins including β-catenin at Ser552.
- Mutating Ser552 to a non-phosphorylatable amino acid in mice resulted in altered CD branching without loss of differentiation in adult CDs.
- The findings point to a role for β-catenin Ser552 phosphorylation in CD branching and sub-segmental CD elongation.

## Introduction

Acting through the V2 receptor (V2R), vasopressin has a variety of structural and functional effects in renal collecting duct principal cells in the mature collecting duct including increased *Aqp2* gene transcription,^1, 2^ increased aquaporin-2 (AQP2) protein half-life,^3, 4^ increased principal cell size,^5–8^ increased exocytosis of AQP2-containing vesicles,^9–11^ decreased AQP2 endocytosis,^10, 12^ increased active Na^+^ reabsorption,^13, 14^ reorganization of actin filaments,^15–19^ and a decreased rate of apoptosis.^20^ Signaling is via the G_α_s/adenylyl-cyclase/cyclic-AMP/protein-kinase-A (PKA) pathway.^21–23^Phosphoproteomic analysis of the effects of vasopressin has identified a number of target proteins that may be responsible for the structural and functional consequences of vasopressin signaling.^24^ Among these targets is β-catenin (Gene symbol: *Ctnnb1*), which undergoes a marked increase in phosphorylation at serine-552 (Ser552) in response to vasopressin, based on both protein mass spectrometry and immunoblotting (**Figure 1**).^25–29^ Here, we investigate the role of Ser552 phosphorylation of β-catenin in the structure of the collecting duct system, using CRISPR-Cas9 to mutate the site to a non-phosphorylatable amino acid.

**Figure 1.**
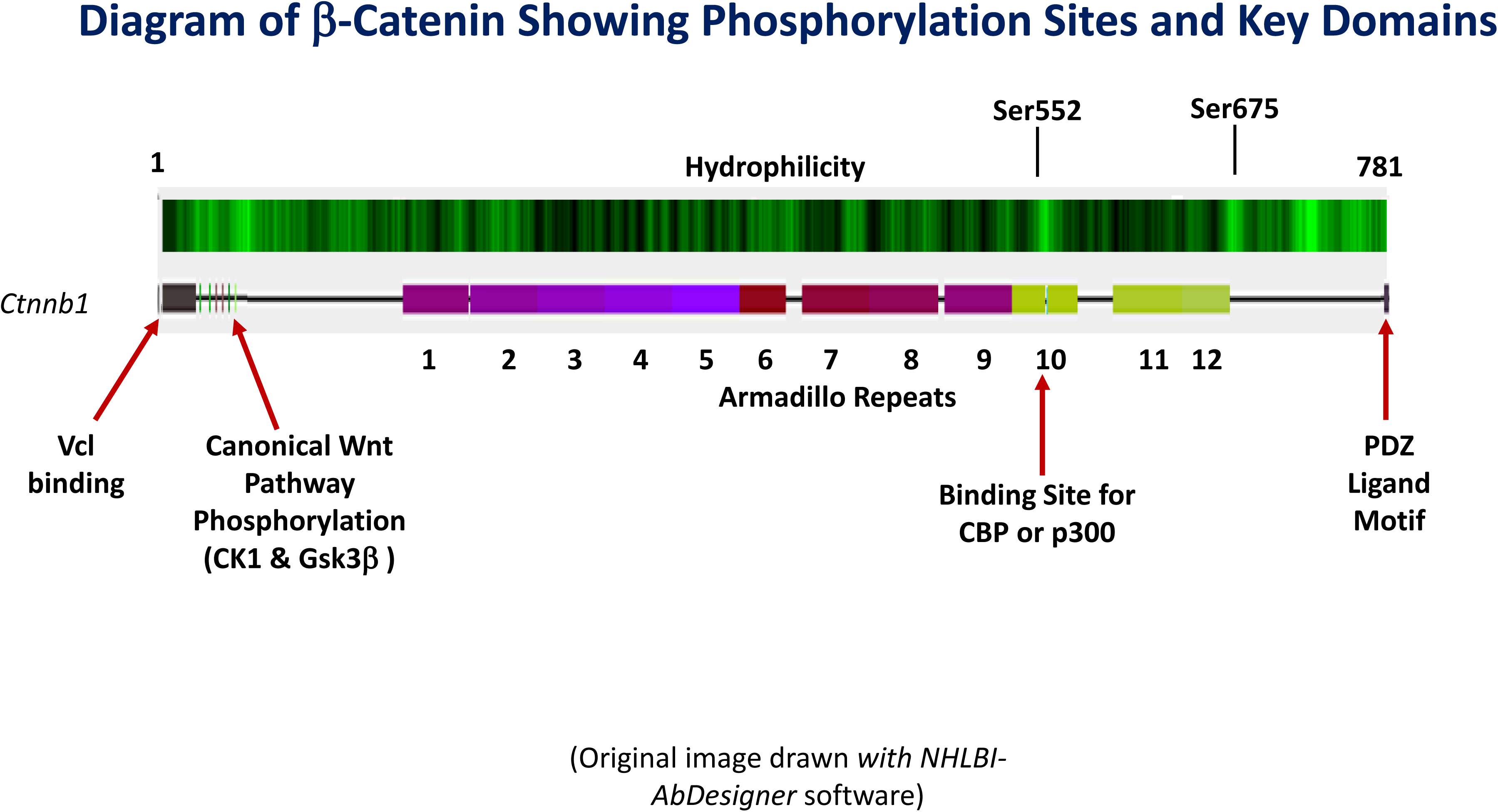
Primary structure of β-catenin showing phosphorylation sites and key domains. The total length in mouse is 781 amino acids (UniProt Accession: Q02248). Original image drawn *with NHLBI-AbDesigner* software (https://esbl.nhlbi.nih.gov/AbDesigner/).^25^ Hydrophilicity calculated using the Kyte-Doolittle algorithm.^26^ Ser552, phosphorylated in response to vasopressin in collecting duct principal cells, lies amid Armadillo Repeat Domain 10, which overlaps the site of binding of histone acetyltransferases CREB-binding protein (CBP) and p300. These histone acetyltransferases regulate chromatin accessibility by increasing histone acetylation, especially at histone H3 lysine-27 (H3K27Ac). Canonical regulation in the Wnt pathway is achieved through phosphorylation of a series of serines near the N-terminus of β-catenin by the protein kinases, casein kinase 1 (CK1) and glycogen-synthase kinase 3 β (GSK3β). Vinculin (Vcl) binding to the N-terminus facilitates β-catenin binding to cell-cell junctions. COOH-terminus interacts with the PDZ protein tax-interacting protein (TIP-1) to inhibit transcription of β-catenin target genes.^27^

Prior studies have implicated the Wnt pathway and β-catenin in the regulation of collecting duct branching during renal development.^30–32^ It is well established that the collecting duct system is derived from the ureteric bud. The detailed morphometric study of Cebrián *et al*.^33^ has tracked the changes in collecting duct architecture from the initial branching event around gestational age 11.5 to birth. Repeated dichotomous branching produces a relatively uniform pattern through gestational age 15.5 with approximately equal distances between branch points.^33^ However, starting at gestational age 15.5, when the outer medulla begins to develop, elongation of the outer medullary portion of the collecting duct occurs to produce the long ‘unbranched segment’, originally observed by Karl Peter^34^ and Jean Oliver.^35^ The detailed and meticulous morphometric studies of Short *et al.*^36^ in mice have demonstrated similar timing and have shown that branching development continues for a few days after birth. Our own extensive studies involving microdissection of cortical collecting ducts (CCDs) from adult rabbits, rats and mice (see for example^37–39^) have shown that the unbranched portion of the collecting duct system extends from the outer medulla into the medullary rays of the cortex. The medullary rays are finger-like projections of the outer medulla into the cortex and are made up of a parallel arrangement of CCDs, cortical thick ascending limbs and S-2 proximal straight tubules. Thus, in the adult kidney, the long unbranched region, consisting of the outer medullary collecting duct (OMCD) and the medullary ray portion of the CCD, separates two branched regions, one in the cortical labyrinth and the other in the inner medulla.

Collecting duct principal cells first become sensitive to vasopressin as they mature perinatally and presumably this is the time when vasopressin-induced phosphorylation of β-catenin at Ser552 first occurs.^40, 41^ That is, vasopressin sensitivity of collecting duct cells coincides with the final stages of branching and formation of the long unbranched collecting duct in the outer medulla and cortical medullary rays. Given the known role of β-catenin in collecting duct development,^30–32^ we hypothesized that mutation of Ser552 to an unphosphorylatable amino acid could alter the ultimate branching architecture of the adult mouse collecting duct system.

## Methods

### Mice

All animal experimental procedures were carried out in accordance with National Heart, Lung, and Blood Institute [NHLBI] animal protoco**l H-0047R6**, approved by the NHLBI Animal Care and Use Committee. Pathogen-free, male, 6-to 8-week-old C57BL/6 mice (Taconic) were used. In order to generate the Ctnnb1 T551A/S552A knock-in mutant mice, CHOPCHOP (https://chopchop.cbu.uib.no) was employed to obtain all the possible sgRNAs, and one sgRNA (CAACGGCGCACCTCCATGGG) was selected and purchased. A single-strand donor oligonucleotide containing the desired mutation was synthesized (IDT, https://www.idtdna.com/pages). The sgRNA (5 ng/ul) and donor oligonucleotides (100 ng/ul) were co-microinjected with Cas9 mRNA (20 ng/ul) into the pronuclei of zygotes collected from C57BL/6 mice. Donor nucleotide sequences are given in **Supplemental Figure 1** (https://esbl.nhlbi.nih.gov/Databases/Catenin-RNA-seq/Data/). Injected embryos were cultured overnight in M16 medium at 37°C with 6% CO₂. Two-cell stage embryos were implanted into the oviducts of pseudo-pregnant surrogate mothers. Offspring were genotyped but overlapping peaks from different mutations in each allele made direct sequencing unclear. To resolve this, PCR products were TOPO-TA-cloned (Thermo Fisher Scientific,#450030), and 6-10 bacterial colonies were sequenced by using QIAprep Spin Miniprep kit (Qiagen, #27106). Subsequent offspring were genotyped by PCR and Sanger sequencing alone. Wild-type (WT) controls were the same C57BL/6 mice.

### Urinary osmolality

Spot urines were collected by holding the mice over a piece of parafilm. Osmolality was determined by vapor pressure osmometry (Wescor).

### Immunoblotting

Immunoblotting was carried out in kidney tissue using standard procedures in our laboratory.^42, 43^ Antibodies are identified in figure legends.

### Microdissection of renal tubule segments

The microdissection of CCDs or branched connecting tubule (CNT)/ initial collecting tubule (iCT) structures from collagenase-treated mouse kidneys was carried out under a Wild M8 dissection stereomicroscope equipped with on-stage cooling as described in the **Supplementary Methods** (https://esbl.nhlbi.nih.gov/Databases/Catenin-RNA-seq/Data/).

### Immunocytochemistry of microdissected CCDs or branched CNT/iCT structures

After the branched CNT/iCT structures or CCDs were isolated, tubules were transferred to a glass bottom culture dish (MatTek, #P35G-0-20-C) coated with Cell-Tak (Corning, # CLS354241-1EA). The tubules were then fixed for 20 minutes with 4% paraformaldehyde in PBS and permeabilized for 20 minutes with permeabilization buffer (PBS with 0.3% Triton X-100 and 0.1% BSA). After permeabilization, tubules were blocked for 30 minutes with blocking buffer (PBS with 1% BSA and 0.2% gelatin). Primary and secondary antibodies are indicated in the figure legends.

### Morphometry of microdissected CCDs

Determination of the numbers of cells per unit length in microdissected CCDs from the mice was carried out using 4’,6-diamidino-2-phenylindole (DAPI) to label nuclei followed by counting of the DAPI-labeled structures (see **Supplemental Methods** for details). The DAPI-labeled microdissected CCDs were also labeled with anti-AQP2 (rabbit, 1:500, Knepper Lab, #K5007) followed by a secondary antibody (Alexa Fluor 568 goat anti-rabbit, 1:400). IMARIS was used to quantify the diameter of the microdissected tubules.

### Glomerular counts in WT and Ser552Ala mice

Whole kidney glomerular counts were done using the acid maceration method of Damidian *et al*.^44^ as modified by Peterson *et al*.^45^ Briefly, a mouse kidney was chopped into small pieces and then reduced to a suspension consisting of glomeruli and small renal tubule fragments by incubation in 6M HCl at 37 °C for 90 minutes. Glomeruli (identified by their round shape) were counted in triplicate in three aliquots of the total suspension under a Wild M8 dissection stereomicroscope and total glomerular number was calculated by multiplying by the ratio of the total volume to the aliquot-volume.

### Preparation of nuclear fractions

Nuclear fractions were prepared from manually dissected inner medullas as described in **Supplementary Methods**.

### Immunocytochemistry of kidney sections

Mice underwent cervical dislocation and were immediately perfused via the left ventricle of the heart with ice-cold DPBS followed by 4% paraformaldehyde in DPBS. The kidneys were post-fixed in 4% paraformaldehyde solution, further processed with alcohol and xylene, and then embedded in paraffin blocks. 5 μm sections were mounted on glass slides for immunofluorescence staining. Dewaxing, antigen retrieval, permeabilization, and antibody labeling are described in **Supplementary Methods**. Immunofluorescent labeling was analyzed using a Zeiss LSM980 confocal microscope using ZENBlue software (Zeiss). The primary and secondary antibodies are listed in the legends.

### Determination of nephron: cortical collecting duct (CCD) ratio in mice

In mice and other mammals, multiple nephrons converge to form a single CCD in the cortical labyrinth via branched CNTs and iCTs.^46^ The ratio of medullary thick ascending limb (MTAL) to outer medullary collecting duct (OMCD) provides a measure of cortical branching ratio (nephrons:CCD) because all structures between the MTAL and OMCD other than the CNT and iCT are unbranched (MTAL, cortical thick ascending limb, CCD in the medullary rays and OMCD).^47^

To measure the MTAL/OMCD ratio, kidneys from WT and Ser552Ala mice underwent perfusion fixation and embedding as described in the previous section. Tissue sectioning was performed to obtain cross-sections of the inner stripe of the outer medulla perpendicular to the corticomedullary axis. The sections underwent immunofluorescence labeling for the Na^+^-K^+^-2Cl^−^ cotransporter, NKCC2 (a marker for the MTAL) and for AQP2 (a marker for OMCD). MTALs and OMCDs were counted in demarcated square fields.

### Small-sample RNA-seq in microdissected CCDs

For RNA-seq analyses, four to eight CCDs were collected for each sample for a total length of 1.5-3.0 mm. cDNA libraries were constructed as described in **Supplementary Methods**. Sequencing was done on an Illumina Novaseq 6000 platform using a 50 bp paired-end modality. FastQC was used to evaluate sequence quality (software version 0.11.9). RNA-seq reads were indexed using STAR (2.7.6a) and aligned to the mouse reference genome from *Ensembl* (release 104) with the matching genome annotation file (release 104).^48^ Transcripts per million (TPM) and expected read counts were generated using RSEM (1.3.3).^49^ Unless otherwise specified, the computational analyses were performed on the NIH Biowulf High-Performance Computing platform.

## Results

### CRISPR-mediated mutation of Ser552 of β-catenin in mouse

To investigate the role of β-catenin phosphorylation at Ser552 in native collecting ducts, we generated a *Ctnnb1* Ser552Ala knock-in mouse using CRISPR/Cas9 (**Figure 2A**). Because the neighboring threonine has sometimes been reported to be phosphorylated,^24^ we also mutated Thr551 to Ala in the same animals, although we will refer to these mice as Ser552Ala knock-in mice in what follows for simplicity. We found that the homozygous mutant mice survived and appeared superficially indistinguishable from WT mice, unlike the global β-catenin-knockout mouse, which was lethal before gastrulation.^50^ The absence of Ser552 phosphorylated β-catenin was confirmed in homozygotes (Ho), and phosphorylation was reduced in heterozygotes (Het) (**Figure 2B**). **Figure 2C** shows immunoblots of the nuclear fractions of the inner medullas of mice showing that the homozygous mutant mice displayed no phosphorylated β-catenin, while total nuclear β-catenin was unaffected (**Figure 2D**). The latter observation demonstrates that the mutation does not prevent nuclear translocation of β-catenin. **Figure 2E** shows random daytime urinary osmolalities for WT and Ser552Ala mice indicating that the Ser552Ala mice are capable of concentrating urine to more than 7 times the plasma level without water restriction or vasopressin administration. **Figure 2F** shows immunofluorescence labeling of ZO-1 and Na^+^-K^+^-ATPase (ATP1A1) in microdissected CCDs in WT and Ser552Ala mice showing no major changes in distribution of these two polarity markers.

**Figure 2.**
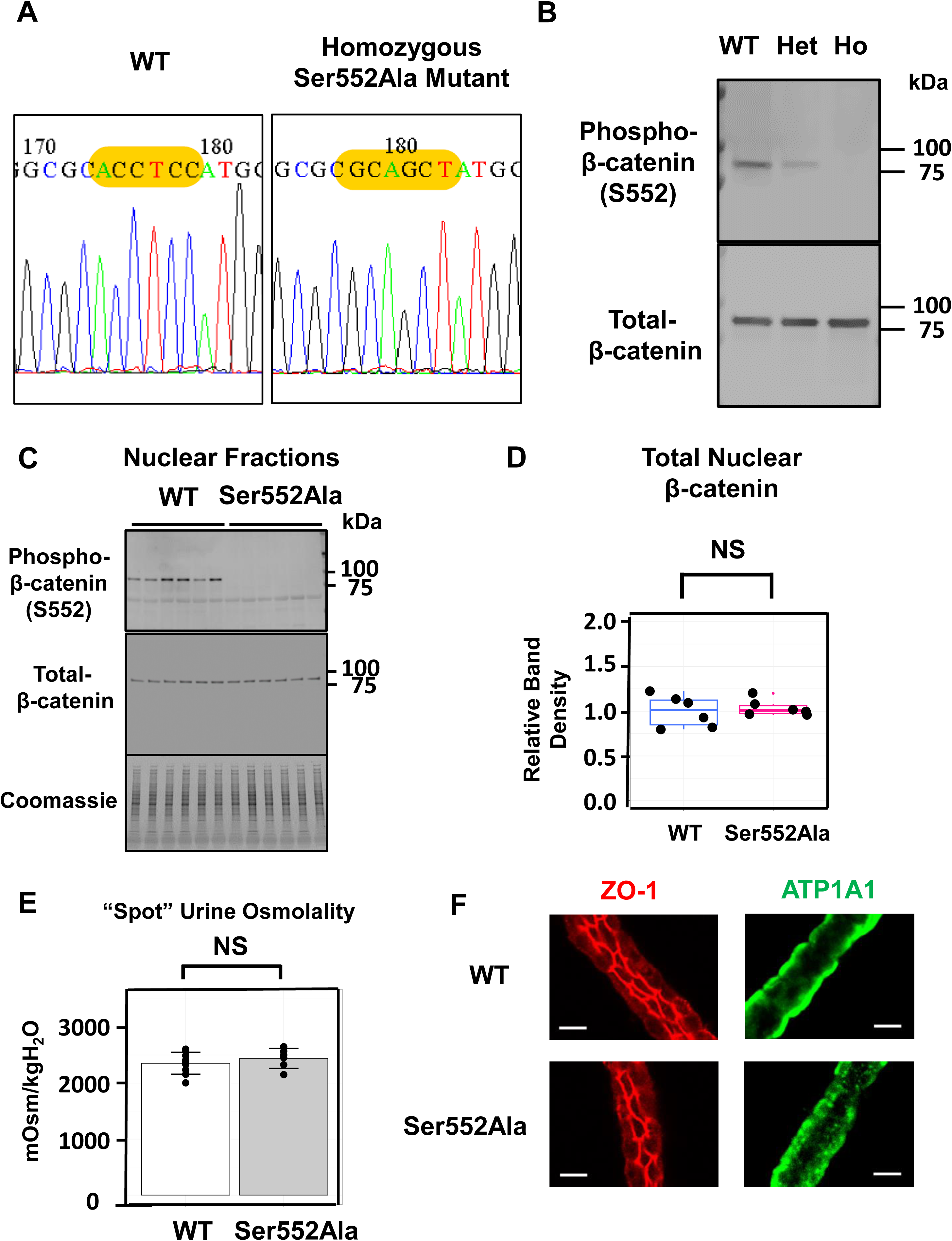
The effects of Ser552Ala versus wild-type (WT) mice on β-catenin phosphorylation, urine osmolality, and epithelial polarity. A. Sanger sequencing demonstrates successful sequence modification in nuclei of Ser552Ala versus WT mice. B. Immunoblotting of whole inner medullas with the phospho-Ser552 antibody shows reduced phosphorylation in heterozygote (Het) and no phosphorylation in homozygote (Ho) with no loss of total β-catenin. C. Immunoblotting of nuclear fractions from inner medullas show no changes in total β-catenin in nuclei of Ser552Ala versus WT mice. Antibodies: anti-CTNNB1 antibody (rabbit, 1:1000, Cell signaling # 9562) or anti-phospho CTNNB1 (Ser552) antibody (rabbit, 1:1000, Cell signaling # 9566). Secondary antibody is goat anti-rabbit IRDye 680 (LI-COR). D. Densitometry of immunoblots shows no change in nuclear abundance of total β-catenin in Ser552Ala mice versus WT mice. NS, not significantly different. E. Urine osmolality in daytime untimed (“spot”) urine collections in Ser552Ala versus WT mice. NS, not significantly different. Urine osmolality was approximately 7 times the plasma osmolality which is _~_300 mOsm/kgH_2_O. F. Lack of effect of Ser552Ala mutation on polarity of microdissected cortical collecting ducts as indicated by labeling of ZO-1 (a tight junction marker) and Na^+^-K^+^-ATPase (ATP1A1, a basolateral membrane marker). Scale bar, 10 µm. Antibodies: anti-ZO-1 (mouse, Invitrogen # 33-9100) and anti-Na^+^/K^+^ ATPase (ATP1A1, rabbit, Knepper Lab, #H7642). The anti-Na^+^/K^+^ ATPase antibody was produced in our laboratory by immunizing rabbits or chickens with a synthetic peptide corresponding to the entire COOH-terminal tail of mouse ATP1A1 protein (sequence C-DEVRKLIIRRRPGGWVEKETYY) conjugated to keyhole limpet hemocyanin (KLH).^66^

A broad survey of 33 tissues and organs, including gross morphology and histology, revealed no major structural abnormalities in the Ser552Ala mice (**Supplementary Spreadsheet** at https://esbl.nhlbi.nih.gov/Databases/Catenin-RNA-seq/Data/). Analysis of blood samples revealed no abnormalities of serum glucose, creatinine, urea, calcium, inorganic phosphorus, total protein, albumin, bilirubin, cholesterol, triglycerides, alkaline phosphatase, creatine kinase, alanine transaminase, aspartate aminotransferase, or lactate dehydrogenase. In addition, there were no abnormalities of any hematologic parameter or blood smear.

### Glomerular counts

Direct counts of the number of glomeruli per kidney showed no differences (13600 ± 1015 per kidney [n=3] in WT versus 13500 ± 889 per kidney [n=3] in Ser552Ala mice, *P* = 0.904), showing that the number of nephrons was unchanged in Ser552Ala mice. This finding fits with the observation that kidney weight normalized to body weight was not different between WT (0.0126±0.0004 [n=3]) and Ser552Ala mice (0.0126±0.0004 [n=3], *P* = 0.979).

### Quantification of cortical tubule branching in WT versus Ser552Ala mice

β-catenin and the Wnt signaling pathway are important determinants of collecting duct branching during development.^30–32^ In the adult mammalian kidney, the endpoint of developmental branching in the cortex are found in CNTs and iCTs, found in the cortical labyrinth. The CNTs and iCTs are the only branched tubule segments prior to the inner medullary collecting ducts (IMCDs).^47^ Branched CNTs and iCTs constitute a manifold-like structure in which several nephrons converge to a single CCD. In rat, on average, slightly greater than 5 nephrons merge to form one CCD^51^ and the nephron:CCD ratio for mouse has been reported to be between 5 and 7.^46^ **Figure 3A** shows a typical cortical branched structure microdissected from a mouse and labeled with antibodies recognizing Na^+^-K^+^-ATPase (ATP1A1) (green), the thiazide sensitive Na^+^-Cl^−^ cotransporter (SLC12A3) (purple) and parvalbumin (PVALB) (cyan). SLC12A3 and PVALB mark multiple distal convoluted tubules that converge into a single CCD.

**Figure 3.**
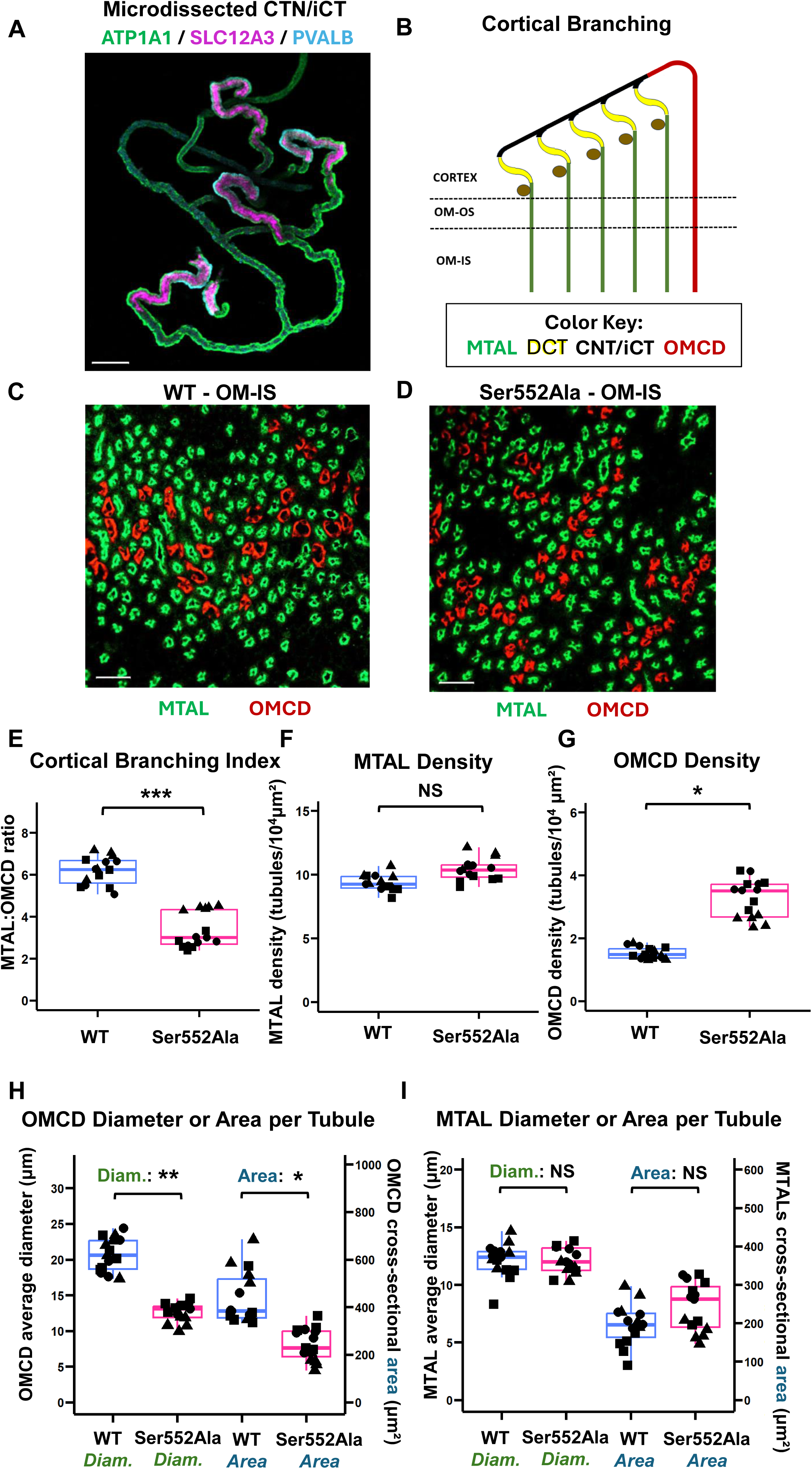
Cortical branching and tubule size in Ser552Ala mice versus wild-type (WT) mice. A. A highly branched cortical structure microdissected from mouse renal cortex showing that distal convoluted tubules converge into a single collecting duct (CCD). Scale bar, 100 µm. Antibodies: anti-Na^+^/K^+^ ATPase (ATP1A1, Millipore, #05-369-25UG), anti-thiazide-sensitive Na^+^-Cl^−^ cotransporter (SLC12A3, Knepper Lab, #4375), and anti-parvalbumin (PVALB, Swant, #GP72). B. Diagram of idealized cortical branching structure demonstrating the approach to quantification of nephron:CCD branching ratio based on counting medullary thick ascending limbs (MTALs, green) and outer medullary collecting ducts (OMCDs, red) in cross sections of mouse inner stripe of outer medulla (OM-IS). C and D. Examples of cross sections of inter stripe of outer medulla labeled for the bumetanide-sensitive Na^+^-K^+^-2Cl^−^ cotransporter (NKCC2, green) and aquaporin-2 (AQP2, red) in WT and Ser552Ala mice, respectively. Scale bar, 50 µm. Antibodies: anti-AQP2 (rabbit, 1:500, Knepper Lab, #K5007) or anti-NKCC2 (chicken, 1:500, Knepper Lab #5198). E. MTAL:OMCD ratio quantification (cortical branching index) in WT versus Ser552Ala mice. ***, *P* < 0.001. F. MTAL count per unit cross sectional area in WT versus Ser552Ala mice. NS, not significantly different. G. OMCD count per unit cross sectional area in WT versus Ser552Ala mice. *, *P* < 0.05. H. Quantification of OMCD diameter and cross-sectional area in WT versus Ser552Ala mice. **, *P* < 0.01; *, *P* < 0.05. I. Quantification of MTAL diameter and cross-sectional area in WT versus Ser552Ala mice. NS, not significantly different.

To measure the cortical branching ratio (nephron:CCD) in WT versus Ser552Ala mice, we used a variant of the technique of Knepper *et al*.^51^ to quantify the MTAL:OMCD ratio in the inner stripe of the outer medulla (ISOM). This ratio is a measure of the average number of cortical branches because the other segments between the MTAL and OMCD are unbranched (**Figure 3B**). To measure the MTAL:OMCD ratio, MTALs and OMCDs were counted in random fields from histological sections taken perpendicular to the corticomedullary axis in the ISOM. We labeled the MTALs using an antibody to the bumetanide-sensitive Na^+^-K^+^-2Cl^−^ cotransporter, NKCC2 (green) and OMCDs with an antibody to AQP2 (red). **Figure 3C and D** show representative fields from WT and Ser552Ala mice, respectively. The measured MTAL:OMCD ratio was significantly greater in the WT mice (6.18 ± 0.66) than in the Ser552Ala mice (3.33 ± 0.82) (**Figure 3E**). This indicates that cortical branching in Ser552Ala mice is attenuated. We asked whether the ratio was different because of a difference in the number of MTALs or OMCDs. The number of MTALs per unit cross sectional area was found to be the same for the two genotypes (**Figure 3F**), consistent with our prior conclusion that the Ser552Ala mice and WT mice have the same number of nephrons based on glomerular counting. In contrast, the number of OMCDs per unit cross sectional area was much higher in the kidneys of Ser552Ala mice (**Figure 3G**). Thus, the difference in branching at the level of the CNTs/iCTs translates to a greater number of collecting ducts in the outer medulla and medullary rays of the cortex. An additional observation (**Figure 3C&D**) is that the OMCDs have smaller diameters in the Ser552Ala mice than in WT. To address this observation quantitatively, we used morphometry of the labeled sections to determine the average diameter and cross-sectional area of MTALs and OMCD (**Figure 3H&I**). As shown in **Figure 3H**, both the diameter and cross-sectional area of OMCDs were significantly lower in Ser552Ala mice. In contrast, there was no difference between WT and Ser552Ala mice with regard to diameter and cross-sectional area of MTALs (**Figure 3I**).

### Effect of Ser552Ala mutation on IMCD

**Figure 4A** shows longitudinal sections of papillae from a WT and a Ser552Ala mouse. Ser552Ala mice had a greater number of collecting ducts with smaller diameters. In separate mice, the papillae were cross-sectioned at approximately 25 µm (upper panels), 75 µm (middle panels), and 125 µm (lower panels) from the papillary tip (**Figure 4B**). **Figure 4C** summarizes total IMCD numbers from the end of the tip. At 25 µm, there was no significant difference in the number of IMCDs between WT (6.33 ± 1.15 [n=3]) and Ser552Ala mice (6.67 ± 0.58 [n=3], *P* = 0.686), indicating that the numbers of duct of Bellini orifices were not likely to be different (**Figure 4C**). However, the number of IMCDs was significantly greater in Ser552Ala mice than in WT mice at a higher level of the papilla (**Figure 4C**).

**Figure 4.**
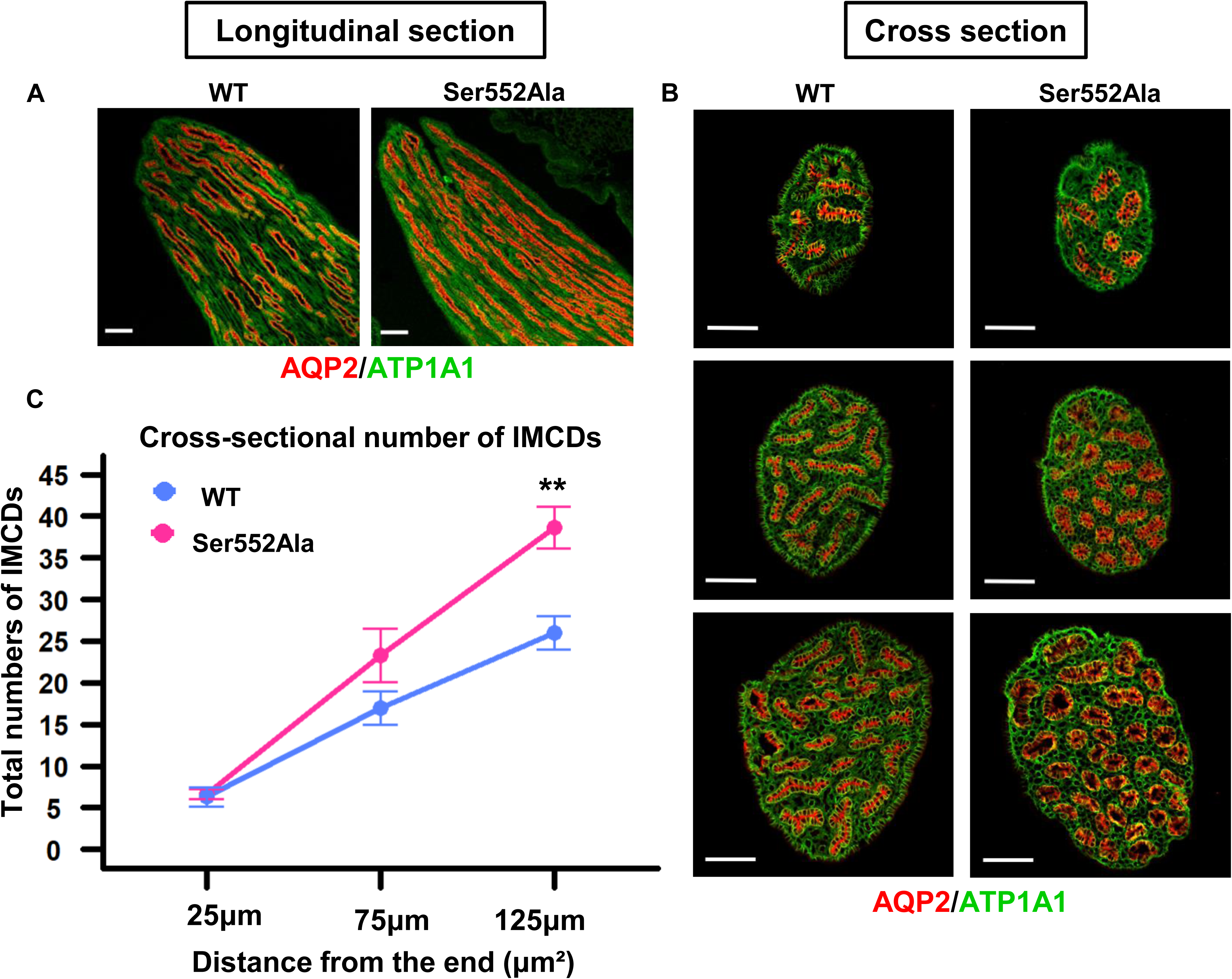
Immunofluorescence-labeled ducts of Bellini in the renal papilla from wild-type (WT) versus Ser552Ala mice. A mouse kidney inner medulla was excised, and the renal papilla was dissected out. The renal papilla was fixed in 4% paraformaldehyde, further processed with alcohol and xylene, and embedded in paraffin blocks. Mouse renal papillae were prepared for either longitudinal or cross-sections and then labeled by immunofluorescence staining. A. Longitudinal section of the renal papilla showed the inner medullary collecting ducts (IMCDs), labeled for aquaporin-2 (AQP2, red) and anti-Na^+^/K^+^ ATPase (green). Scale bar, 100 µm. B. Cross-section of the renal papilla showing the ducts of Bellini at the renal papillary tip (upper panel: within 25 µm from the end) and their increased numbers at higher levels (middle panel: 75 µm, lower panel: 125 µm from the end). Tissue labeled for AQP2 (red) and anti-Na^+^/K^+^ ATPase (green). Scale bar, 100 µm. Antibodies: anti-AQP2 (rabbit, 1:500, Knepper Lab, #K5007), anti-Na^+^/K^+^ ATPase (mouse, 1:400, Millipore, #05-369-25UG). C. The total numbers of IMCDs at different distances of the renal papillary tip (25 µm, 75 µm, or 125 µm from the end) in WT (blue) versus Ser552Ala mice (pink). **, *P* < 0.01.

### Effect of Ser552Ala mutation on the CCD transcriptome

Since β-catenin functions as a transcriptional co-regulator, we asked whether CCDs from adult Ser552Ala mice manifest changes in gene expression as measured by RNA-seq. We carried out RNA-seq in microdissected CCDs from Ser552Ala and WT mice that were otherwise untreated. The CCD samples were analyzed for each of four wild-type mice (n=4) and each of four Ser552Ala mice (n=4) (Figure 5). The full curated data set is available at https://esbl.nhlbi.nih.gov/Databases/Catenin-RNA-seq/.

**Figure 5.**
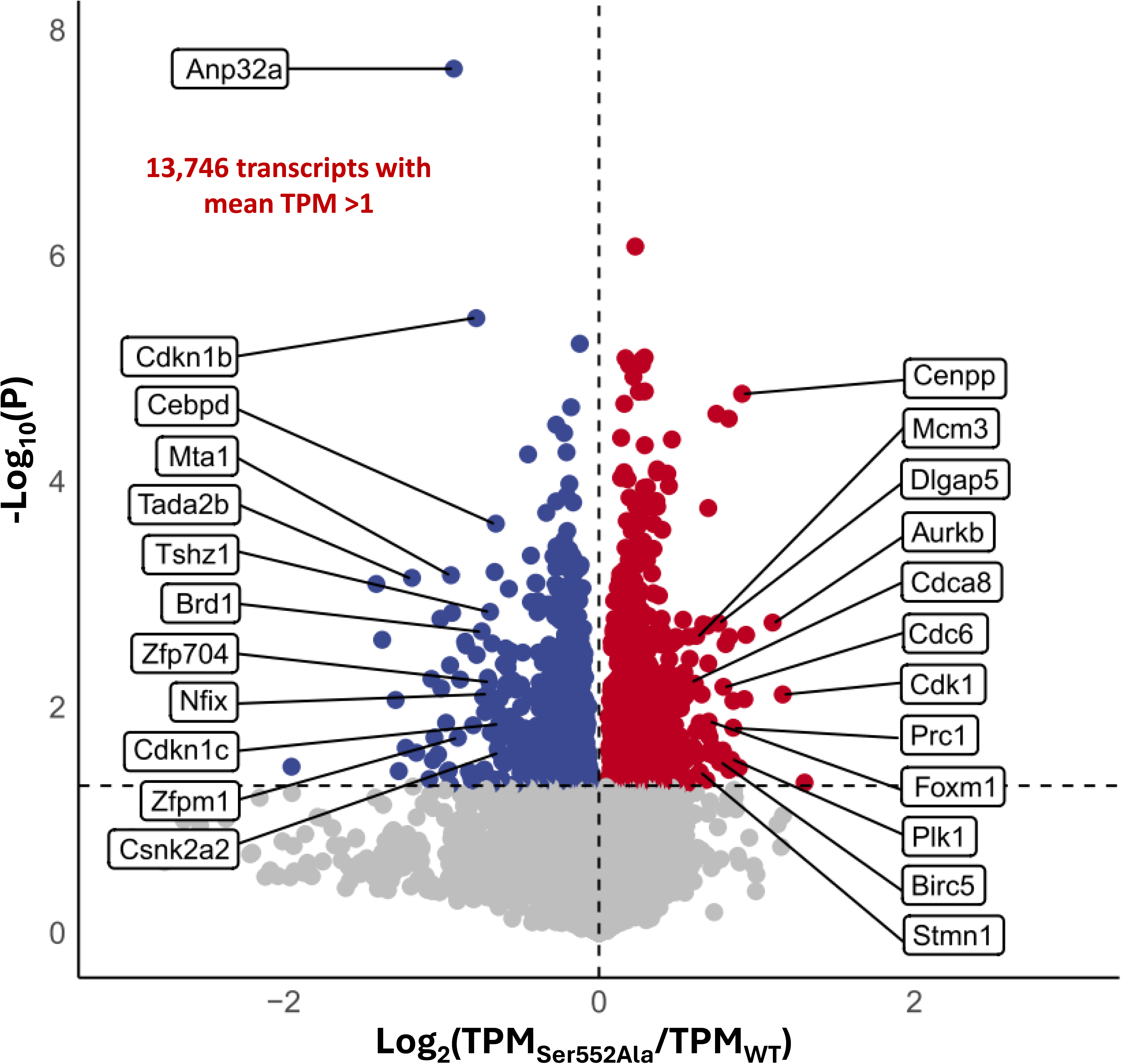
Volcano plot representation of transcriptomic changes in microdissected mouse cortical collecting ducts (CCDs) from wild-type (WT) versus Ser552Ala mice. TPM, transcripts per million. *P* is derived from the t-statistic in a comparison of four Ser552Ala mice versus four WT mice. Three technical replicates (separate microdissected CCDs) were used to calculate a mean value for each biological replicate (animal). The total number of biological replicates was n=4 for each genotype. Gene symbols on the right indicate increased transcripts that match the *Gene Ontology Biological Process* term “mitotic cell cycle process”. Downregulated transcripts on the left indicated decreased transcripts that map to the *Gene Ontology Biological Process* term “animal organ development”. Out of 13,746 transcripts with mean TPM greater than 1 in WT, 42 were identified as increased in Ser552Ala mutant CCDs and 64 were identified as decreased based on dual criteria (|log_2_[Ser552Ala/WT] |> 0.60 and *P* <0.05). (The threshold of 0.60 was calculated from the entire population of log_2_[Ser552Ala/WT] values to approximate the 95% confidence interval using the Empirical Bayes’ Method.^52^). Raw data was deposited at GEO (https://www.ncbi.nlm.nih.gov/geo/query/acc.cgi?acc=GSE291074, Secure token: yvaresyoxrmlfaf).

Out of 13,746 transcripts with mean TPM greater than 1 in WT, 42 were identified as increased in Ser552Ala mutant CCDs and 64 were identified as decreased based on dual criteria (|log_2_[Ser552Ala/WT] |> 0.60 and *P* <0.05)^52^. Gene-set enrichment analysis revealed that upregulated transcripts mapped highly significantly to *Gene Ontology Biological Process* terms related to cell division (*P* < 10^−6^) including “mitotic cell cycle process”, “mitotic cytokinesis”, “nuclear division”, “G2/M transition of mitotic cell cycle”, and “regulation of cell cycle process” (**Table 1A**). Control analysis using 42 randomly selected genes revealed no *Biological Process* terms with Fisher Exact P values less than 1.3×10^−2^. Transcripts associated with the *Gene Ontology Biological Process* term “mitotic cell cycle process” are labeled on the right-hand side of Figure 5. Among the “mitotic cell cycle process” gene set elements are four key protein kinases involved in cell-cycle regulation and mitosis: aurora kinase B (*Aurkb*), cyclin-dependent kinase 1 (*Cdk1*), NIMA-related expressed kinase 5 (*Nek5*), and polo-like kinase 1 (*Plk1*). Interestingly, one of the upregulated transcripts codes for the transcription factor *Foxm1* (also known as “*Trident*”), a key regulator of the expression of cell cycle genes essential for DNA replication and mitosis.^53–55^

**Table 1.** *Gene Ontology Biological Process* terms over-represented in microdissected cortical collecting ducts of Ser552Ala versus wild-type mice based on (A) upregulated transcripts and (B) downregulated transcripts.

**A. Upregulated Transcripts**
| Term | % of Total | *P*<br>(Fisher Exact) | Fold Enrichment | Transcripts |
| --- | --- | --- | --- | --- |
| mitotic cell cycle process | 37.5 | $8.2 \times 10^{-11}$ | 7.91 | Plk1,Cdca8,Kif22,Cdc6,Foxm1,Aurkb, Ncaph,Anln,Prc1,Stmn1,Cdk1,Mcm3,Birc5, Ect2,Dlgap5 |
| mitotic cytokinesis | 17.5 | $8.8 \times 10^{-10}$ | 34.63 | Anln,Plk1,Stmn1,Cdca8,Birc5,Ect2,Aurkb |
| nuclear division | 27.5 | $9.3 \times 10^{-9}$ | 9.40 | Top2A,Prc1,Plk1,Stmn1,Cdca8,Birc5,Kif22, Cdc6,Dlgap5,Ncaph,Aurkb |
| spindle organization | 17.5 | $6.3 \times 10^{-7}$ | 13.52 | Prc1,Plk1,Stmn1,Cdca8,Birc5,Dlgap5, Aurkb |
| G2/M transition of mitotic cell cycle | 15 | $8.3 \times 10^{-7}$ | 17.97 | Plk1,Cdk1,Birc5,Cdc6,Foxm1,Aurkb |
| regulation of cell cycle process | 22.5 | $2.3 \times 10^{-5}$ | 5.48 | Prc1,Plk1,Cdk1,Cdca8,Birc5,Cdc6,Ect2, Ncaph,Aurkb |
| microtubule cyto-skeleton organization | 17.5 | $5.8 \times 10^{-4}$ | 4.67 | Prc1,Plk1,Stmn1,Cdca8,Birc5,Dlgap5, Aurkb |
| DNA metabolic process | 20 | $3.6 \times 10^{-3}$ | 3.04 | Top2a,Cdk1,Mcm3,Cyp1b1,Kif22,Cdc6, Foxm1,Aurkb |
| apoptotic process | 25 | $7.2 \times 10^{-3}$ | 2.34 | Top2a,Plk1,Ifi203,Cdk1,Cyp1b1,Birc5, S100a14,Ect2,Tlr4,Aurkb |

**B. Downregulated Transcripts**
| Term | % of Total | *P*<br>(Fisher Exact) | Fold Enrichment | Transcripts |
| --- | --- | --- | --- | --- |
| animal organ development | 28.33 | $6.5 \times 10^{-3}$ | 1.81 | Cdkn1c,Egln1,Ctla2a,Brd1,Cdkn1b,Nfix,Cebpd, Tshz1,Cul3,Csnk2a2,Ush1c,2810429I04RIK, Ajap1,Oc90,H4c18,Gcnt2,Zfpm1 |

\* Calculations carried out using *Database for Annotation, Visualization and Integrated Discovery* (DAVID) at https://david.ncifcrf.gov/. Background gene list included all transcripts with mean TPM>1.

Gene-set enrichment analysis with the downregulated transcripts mapped significantly to only one *Gene Ontology Biological Process* term: “animal organ development” (**Table 1B**) (labeled on left-hand side of Figure 5). Control analysis using 64 randomly selected genes revealed no *Biological Process* terms with Fisher Exact *P* values less than 5.8×10^−3^. Among the downregulated transcripts in the CCDs of Ser552Ala mice were two cyclin-dependent kinase inhibitor proteins (*Cdkn1b* and *Cdk1c*), which are negative regulators of cell cycle progression and proliferation.^56, 57^ Aside from *Cdkn1b* and *Cdkn1c* mRNAs, a number of transcripts that code for proteins involved in regulation of transcription were decreased in abundance, including transcription factors (*Cebpd*, *Nfix*, *Zfp704*), transcriptional coregulators (*Mta1*, *Tada2b*, *Tshz1*, *Zfpm1*), regulators of histone acetylation (*Anp32a* and *Brd1*), and a protein kinase with a key role in Wnt/β-catenin signaling (*Csnk2a2*) (Figure 5).

Despite the induction of several transcripts involved in the cell cycle, there was no evidence of dedifferentiation as shown in **Table 2**, indicating that the abundances of receptors, water channels, potassium channels and sodium channels characteristic of differentiated collecting duct principal cells were unaffected by the Ser552Ala mutation. Notably, the mutation did not significantly affect the abundance of the transcript coding for AQP2.

**Table 2.** Transcripts corresponding to collecting duct principal cell marker genes: RNA-seq in microdissected cortical collecting ducts of Ser552Ala versus wild-type mice.

| Gene Symbol | Annotation | Wild-type mean TPM | Mutant mean TPM | log<sub>2</sub> (Mutant/Wildtype) | P |
| --- | --- | --- | --- | --- | --- |
| **Acer2** | alkaline ceramidase 2 | 50.2 | 52.7 | 0.07 | 0.597 |
| **Aqp2** | aquaporin 2 | 17256.9 | 15649.1 | -0.14 | 0.497 |
| **Aqp3** | aquaporin 3 | 4388.3 | 4502.5 | 0.04 | 0.721 |
| **Aqp4** | aquaporin 4 | 173.2 | 198.0 | 0.19 | 0.688 |
| **Atp1a1** | ATPase, Na<sup>+</sup>/K<sup>+</sup> transporting, alpha 1 polypeptide | 450.7 | 397.6 | -0.18 | 0.264 |
| **Atp1b1** | ATPase, Na<sup>+</sup>/K<sup>+</sup> transporting, beta 1 polypeptide | 1179.5 | 1230.9 | 0.06 | 0.102 |
| **Avpr2** | arginine vasopressin receptor 2 | 1032.7 | 1055.2 | 0.03 | 0.836 |
| **Clcnkb** | chloride channel, voltage-sensitive Kb | 1619.7 | 1615.2 | 0.00 | 0.959 |
| **Fxyd2** | FXYP domain-containing ion transport regulator 2 | 981.6 | 747.8 | -0.39 | 0.168 |
| **Fxyd4** | FXYP domain-containing ion transport regulator 4 | 5855.2 | 6513.7 | 0.15 | 0.413 |
| **Fzd1** | frizzled class receptor 1 | 45.2 | 47.0 | 0.06 | 0.675 |
| **Gata3** | GATA binding protein 3 | 126.6 | 122.6 | -0.05 | 0.575 |
| **Kcnj1** | potassium inwardly-rectifying channel, subfamily J, member 1 | 987.6 | 1050.6 | 0.09 | 0.464 |
| **Lgals3** | lectin, galactose binding, soluble 3 | 1616.4 | 1667.3 | 0.04 | 0.419 |
| **Lpar1** | lysophosphatidic acid receptor 1 | 9.3 | 11.0 | 0.24 | 0.110 |
| **Npy6r** | neuropeptide Y receptor Y6 | 31.4 | 34.3 | 0.13 | 0.516 |
| **Nr3c2** | mineralocorticoid receptor | 85.5 | 77.9 | -0.13 | 0.044 |
| **Ptger4** | prostaglandin E receptor 4 (subtype EP4) | 21.3 | 20.8 | -0.03 | 0.832 |
| **Ptges** | prostaglandin E synthase | 260.2 | 242.9 | -0.10 | 0.380 |
| **Scnn1a** | sodium channel, nonvoltage-gated 1 alpha | 396.8 | 434.3 | 0.13 | 0.066 |
| **Scnn1b** | sodium channel, nonvoltage-gated 1 beta | 1079.6 | 1113.0 | 0.04 | 0.368 |
| **Scnn1g** | sodium channel, nonvoltage-gated 1 gamma | 777.9 | 809.3 | 0.06 | 0.247 |
| **Stc1** | stanniocalcin 1 | 57.2 | 63.5 | 0.15 | 0.377 |
| **Tmem45b** | transmembrane protein 45b | 668.0 | 666.6 | 0.00 | 0.968 |
| **Tspan1** | tetraspanin 1 | 633.1 | 633.7 | 0.00 | 0.993 |

\* Collecting duct principal cell marker genes from https://esbl.nhlbi.nih.gov/Signaling-Pathways/PC-Diff-Markers/
Abbreviations: TPM, transcripts per million; P, P value from t-statistic.

Although the single-tubule RNA-seq in CCDs demonstrated a pattern of expression changes that point to a possible role for β-catenin phosphorylation at Ser552 to inhibit cellular proliferation, it is not clear whether these transcriptional changes are accompanied by overt cell proliferation or alternatively are associated with entry into the cell cycle with cell-cycle arrest. To address these possibilities in the CCD, we carried out DAPI labeling of microdissected CCDs from Ser552Ala mice and WT mice (Figure 6A). The number of cells per unit length determined by counting DAPI-labeled nuclei was not significantly different between WT and Ser552Ala mice (Figure 6B). However, as seen previously for OMCDs, the diameters of CCDs from Ser552Ala mice were significantly smaller than in WT mice (Figure 6C). DNA content in nuclei can be assessed by measuring DAPI fluorescence intensity integrated over the entire nucleus. We expect nuclear DNA content to increase in S phase and to be maintained in G2 phase and early M phase. Figure 6D shows a histogram of nuclear DAPI levels. The mean per-nucleus DAPI fluorescence is higher in Ser552Ala mice compared to WT (162.3 ± 46.8 [n=4] vs 150.8 ± 42.6 [n=4], *P* = 0.005). Nuclei with total DAPI fluorescence greater than the indicated threshold (mean plus one standard deviation of total fluorescence) were classified as “high-intensity” nuclei. The proportion of cells with high-intensity nuclei was significantly higher in Ser552Ala mice compared to WT (26.7% *vs*. 13.8%, *P* = 0.0005, Fisher exact test). Overall, the DAPI fluorescence data are consistent with a lack of overt cellular proliferation, but with a higher frequency of cells arrested in G2/M.

**Figure 6.**
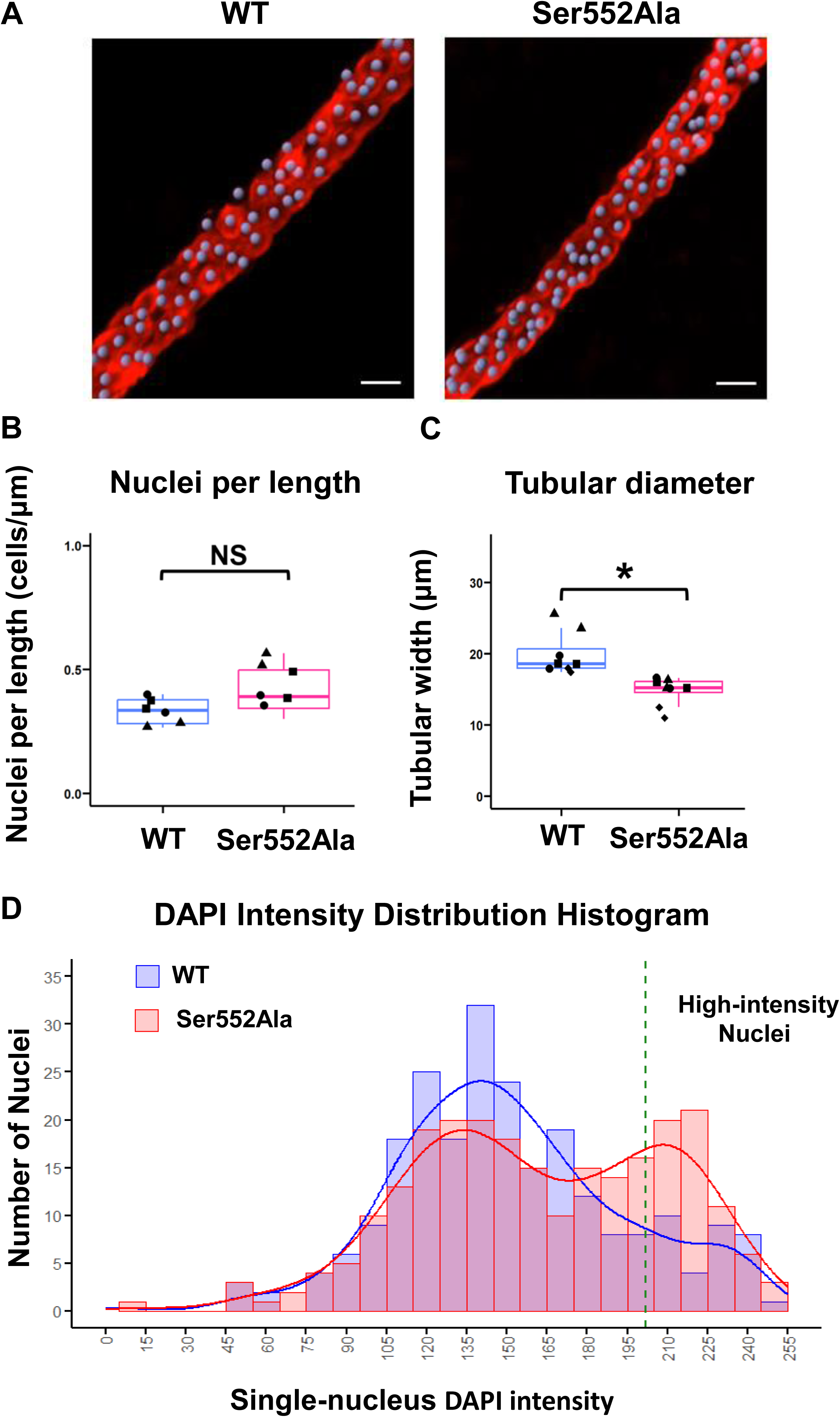
Nuclear counting and cell cycle indices in immunofluorescence-labeled microdissected cortical collecting ducts (CCDs) from wild-type (WT) versus Ser552Ala mice. After microdissection, CCDs were fixed with 4% paraformaldehyde, followed by labeling for aquaporin-2 (AQP2) and for nuclei using 4’,6-diamidino-2-phenylindole (DAPI). A. IMARIS-generated confocal immunofluorescent analytical image of microdissected CCDs from WT and Ser552Ala mice. AQP2 is labeled in red. Nuclei are labeled with DAPI and shown in grayish purple based on IMARIS “spot” analysis tools. Scale bar, 10 µm. Antibodies: anti-AQP2 (rabbit, 1:500, Knepper Lab, #K5007). B and C. Nuclei per length and tubular diameter of microdissected CCDs in WT versus Ser552Ala mice. *, *P* < 0.05; NS, not significantly different. D. Histogram of DAPI immunofluorescence intensity in nuclei from microdissected CCDs of WT and Ser552Ala mice. The x-axis represents DAPI-labeled single-nucleus intensity, and the y-axis represents the number of nuclei. The blue and pink bars indicate the number of nuclei for WT and mutant mice, respectively, with overlaid density curves for each group. A green dashed line marks the threshold for “high-intensity” nuclei (defined as the mean plus one standard deviation of total immunofluorescence).

## Discussion

In the collecting duct system of adult mouse kidney, branching is seen only in the cortical labyrinth and inner medulla, but not in the collecting ducts of the cortical medullary rays (CCDs) or outer medulla (OMCDs) (**Introduction**). The CCDs and OMCDs are long unbranched segments that form by inter-branch elongation starting at gestational age 15.5.^33^ Our data indicate that, in total, there are 11 generations of dichotomous branches (calculated as log_2_[13600/6.3] from Figure 7) in the collecting duct system of WT mice, consistent with the findings of Cebrián *et al*.^33^ The β-catenin Ser552Ala mice also have 11 generations of dichotomous branches, but they are divided differently between the cortical labyrinth (fewer in mutant) and inner medulla (more in mutant). Within the branching architecture, the long unbranched OMCD/CCD occurs at average generation level 8.4 (log_2_[2200/6.3] from Figure 7) in the WT mice and at average generation level 9.2 (log_2_[4100/6.7]) in the Ser552Ala mice. Because elongation occurs higher up in the branched tree of the collecting duct system, there are more OMCDs and CCDs in the Ser552Ala mice than in WT mice, which if other factors are equal would predict more overall transport of sodium, potassium and water. However, we found that these collecting ducts have, on average, smaller diameters. The smaller diameters may compensate for the larger collecting duct counts with regard to total luminal surface area available for transport.

**Figure 7.**
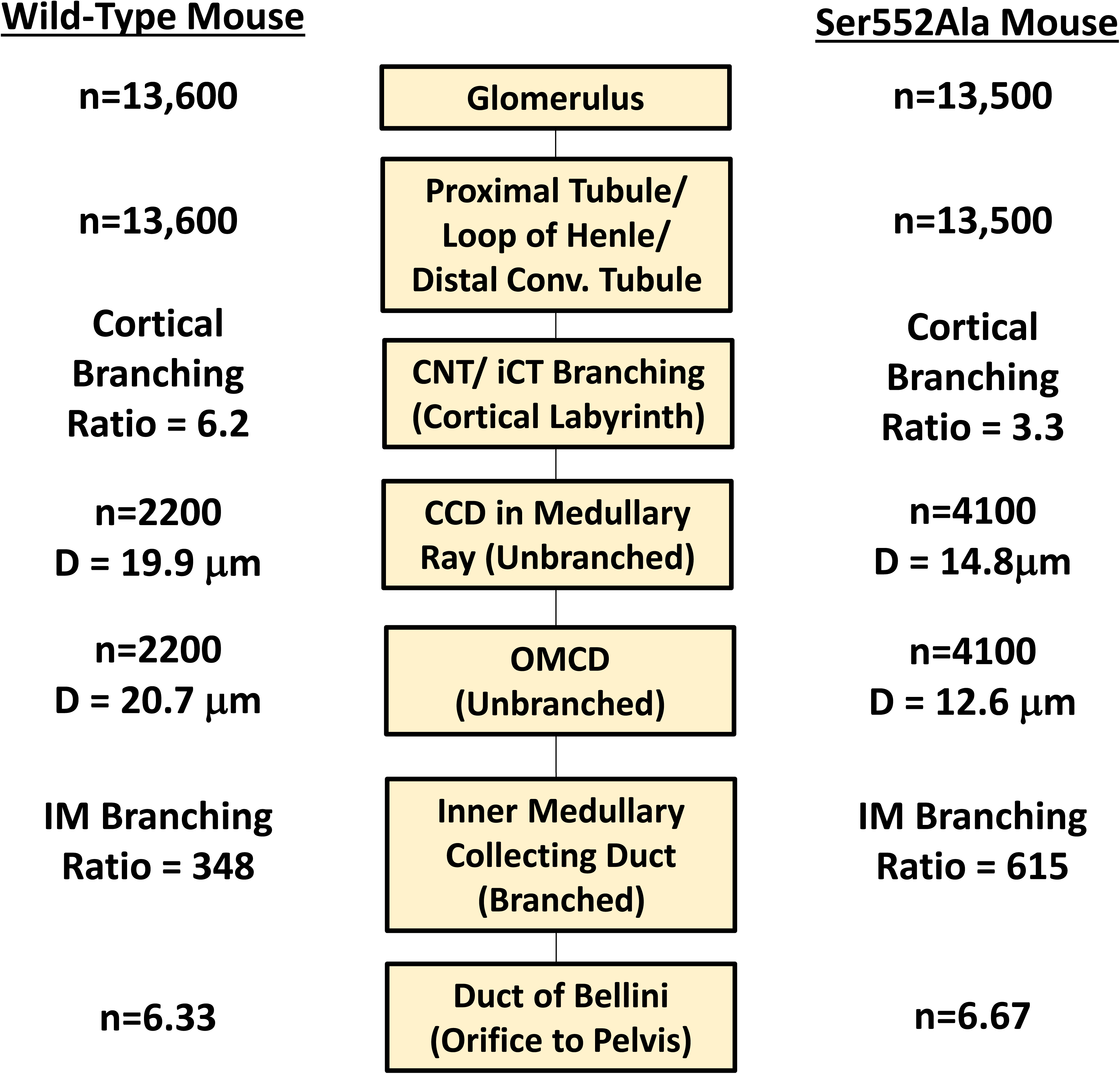
Schematic diagram summarizing structural parameters in wild-type (WT) and Ser552Ala mice. The diagram provides a visual representation of tubule branching patterns, highlighting the structural organization and differences in branching ratios in WT and Ser552Ala mice. Glomerular counts show approximately equal nephron numbers (n = 13,600 in WT mice, n = 13,500 in Ser552Ala mice). Distal convoluted tubules connect to connecting tubules (CNT) which undergo convergence through branched CNTs and initial collecting tubules (iCTs) and then connect to a cortical collecting duct (CCD). The medullary thick ascending limb (MTAL)/outer medullary collecting duct (OMCD) ratio, which quantifies the numbers of nephrons converging into an unbranched collecting duct, averaged 6.2 in WT mice and 3.3 in Ser552Ala mice. The unbranched collecting ducts (CCD/OMCD) are more numerous in the mutant mice (n = 2,200 in WT mice, n = 4,100 in Ser552Ala mice). The convergence of collecting ducts resumes in the inner medulla, eventually reducing inner medullary collecting duct (IMCD) number (ducts of Bellini) to 6.33 in WT mice and 6.67 in Ser552Ala mice.

To address the mechanisms that govern the timing between collecting duct branching and elongation, extensive additional studies will be needed to address two questions: 1) What are the signals that are responsible for local elongation of collecting ducts in the developing outer medulla and cortical medullary rays?; and 2) How is the rate of branching controlled and eventually stopped in the post-natal period? As reviewed in the **Introduction**, collecting ducts first acquire vasopressin sensitivity in the perinatal period, which overlaps the final stages of collecting duct branching and collecting duct elongation. Thus, vasopressin signaling, involving β-catenin phosphorylation, is quite possibly a determinant of collecting duct branching architecture through effects on collecting duct branching and/or elongation. Wnt signaling via *Wnt7b* and β-catenin are important for elongation of the renal medulla as a whole including the loop of Henle,^58^ and we note that vasopressin also markedly increases β-catenin phosphorylation at Ser552 in the medullary thick ascending limb of rat.^59^

The effect of the β-catenin Ser552Ala mutation on gene expression in adult CCDs (RNA-seq) is relatively subtle, with selective changes in transcripts involved in regulation of cell cycle and organ development (**Table 1**). However, cell counts per unit length of collecting ducts showed no significant changes. The increase in mean DNA content and in the number of high DNA-content cells (Figure 6D) is consistent with the possibility that there are more collecting duct cells in the cell cycle, but with G2/M arrest rather than frank proliferation. Among the transcripts decreased in abundance in the mutant CCDs were two cyclin-dependent kinase inhibitor proteins (*Cdkn1b* and *Cdk1c*), which are negative regulators of cell cycle progression and proliferation.^56, 57^ *Cdkn1b* is known as p27 (or Kip1), while *Cdkn1c* is known as p57 (or Kip2). They both work by inhibiting the cyclin E-CDK2 and cyclin D-CDK4/6 complexes. *Cdkn1b*/p27 is involved in regulation of cell ‘quiescence’.^60^ Cell quiescence is a reversible state in which cells exit the cell cycle (G0 phase) and maintain functional differentiation. Quiescent cells, unlike senescent cells, can be induced to re-enter the cell cycle and resume proliferation.^61^ *Cdkn1c*/p57 has a special role in development, and is essential for proper organogenesis.^62–64^

A topic for future studies is the mechanism by which β-catenin phosphorylation at Ser552 may alter gene expression. A clue is offered by the data of Takemaru and Moon^65^ showing that Armadillo domain 10, which contains Ser552, is the site of p300 or CREB-binding protein (CBP) binding to β-catenin (Figure 1). CBP and p300 are histone acetyltransferases that alter chromatin structure and increase local DNA accessibility. Thus, it appears possible that transcription may be altered by Ser552 phosphorylation as a consequence of altered CBP- or p300-mediated changes in the chromatin structure surrounding target genes.

## ACKNOWLEDGEMENTS

The authors thank Dr. Chengyu Liu and Dr. Fan Zhang (NHLBI Transgenic Core) for creating the mutant mice. Next-generation sequencing was done in the NHLBI DNA Sequencing Core Facility (Yuesheng Li, Director). Light microscopy images were taken in the NHLBI Confocal Microscopy Core Facility (Dr. Christian Combs, Director). Dr. Matthew Starost carried out pathological surveys of mouse phenotype. Computational analyses were performed on the NIH Biowulf High-Performance Computing platform.

## DATA SHARING

Curated RNA-seq data are accessible at https://esbl.nhlbi.nih.gov/Databases/Catenin-RNA-seq/.

Sequencing data (FASTQ) have been deposited in the GEO: https://www.ncbi.nlm.nih.gov/geo/query/acc.cgi?acc=GSE291074 (Secure token: yvaresyoxrmlfaf).

## SUPPLEMENTAL MATERIAL

**Supplementary Spreadsheet. Pathological survey and blood hematology and biochemistry analysis of wild-type and Ser552Ala mice.**

**Supplementary Figure 1. Donor oligonucleotide sequences used for CRISPR/Cas9-mediated generation of Ser552Ala mice.**

**Supplementary Methods. Detailed descriptions of experimental procedures and methods used in this study.**

**Available for download at:** https://esbl.nhlbi.nih.gov/Databases/Catenin-RNA-seq/Data/

